# A validation of Illumina EPIC array system with bisulfite-based amplicon sequencing

**DOI:** 10.1101/2020.05.25.115428

**Authors:** Alexandra Noble, John Pearson, Joseph Boden, John Horwood, Neil Gemmell, Martin Kennedy, Amy Osborne

## Abstract

The Illumina Infinium® MethylationEPIC BeadChip system (hereafter EPIC array) is considered to be the current gold standard detection method for assessing DNA methylation at the genome-wide level. EPIC arrays are used for hypothesis generation or pilot studies, the natural conclusion is to validate methylation candidates and expand these in a larger cohort, in a targeted manner. As such, an accurate smaller-scale, targeted technique, that generates data at the individual CpG level that is equivalent to the EPIC array, is needed. Here, we tested an alternative DNA methylation detection technique, known as bisulfite-based amplicon sequencing (BSAS), to determine its ability to validate CpG sites detected in EPIC array studies. BSAS was able to detect differential DNA methylation at CpG sites to a degree which correlates highly with the EPIC array system. However, BSAS correlated less well with EPIC array data when the magnitude of change via EPIC array was greater than 5%, suggesting that this lower specificity at larger differential methylation values is a consequence of PCR amplification that BSAS requires. However, our data suggests that BSAS does offer advantages that the EPIC array: BSAS amplifies a region of the genome (~500bp) around a CpG of interest, allowing analyses of other CpGs in the region that may not be present on the EPIC array, aiding discovery of novel CpG sites and differentially methylated regions of interest. We conclude that BSAS offers a valid investigative tool for specific regions of the genome that are currently not contained on the array system.

## Introduction

Epigenetic modifications, such as DNA methylation, play a vital role in regulating gene expression [1] and have the potential to induce phenotypic changes [2–6]. DNA methylation occurs when a methyl group is covalently transferred to the C5 position of the cytosine ring of a DNA molecule by a methyltransferase enzyme, with the resulting modified cytosine then termed 5-methylcytosine (5mC) [7]. In mammals, most DNA methylation occurs at CpG dinucleotides. CpG sites themselves can be defined as a singular modified cytosine residue which reside predominantly in promoter regions of the genome, which are renowned for being CpG dense [8].

DNA methylation is heavily influenced by the surrounding environment; factors such as tobacco smoking [9–12], alcohol [13, 14], nutrition [15, 16], stress [17] and aging [18, 19] can all impact on DNA methylation at CpG sites. Alterations to DNA methylation are associated with changes in phenotype and also, in some instances, methylation changes contribute to disease pathology [20–23].

As a result of these relatively recent observations, the assessment of differential DNA methylation in humans, and in particular, epigenome-wide association studies (EWAS), is a burgeoning field. High-throughput array technologies are a popular choice for EWAS, due to their robustness and accuracy [24]. The Illumina Infinium^®^ MethylationEPIC array (hereafter ‘EPIC array’) quantifies methylation at 850,000 different CpG sites [25], and although this is still a small proportion of the total number of CpG sites in the genome (~28 million [26]) it represents a broad distribution of sites that give a specific and robust measurement of methylation at those sites.

Further, the goal of many whole-genome studies of DNA methylation is often a pilot or scoping study to capture a range of targets that may be associating with, e.g., a particular environmental exposure. As such, once the genome has been investigated in a number of samples, a whole-genome approach is not always necessary if the user simply requires follow up and/or validation of identified loci in a larger cohort. This means that accurate normalisation of raw data is imperative to ensuring data is robust for follow-up analysis and/or validation. To undertake further analyses and to validate methylation array-based experiments, several different methods exist that that rely on bisulfite treatment of DNA: bisulfite-based amplicon sequencing (BSAS), bisulfite pyrosequencing and methylation-specific PCR (MS-PCR) are methods which can specifically target a predetermined area of interest in the genome at a low cost and higher sample throughput, compared to arrays. Bisulfite pyrosequencing has been found to show congruence to EPIC array analysis [27]. However, pyrosequencing technology is known to have quantitative flaws due to the output of sequences generated through luminescent methods [28]. MS-PCR is a method often used in clinical settings [29], however it has a high false positive rate [30]. By contrast, BSAS detects cytosine methylation to base-pair scale resolution without reliance on light detection methods for sequencing [31]. BSAS is a multiplex procedure that can quantitively assess each CpG site within numerous target regions at the same time [32]. Thus, given the limitations of pyrosequencing and MS-PCR, here we examine whether BSAS is an accurate EPIC array validation, replication and/or expansion tool for targeted DNA methylation analyses.

To answer the question, we used data from EPIC arrays conducted on individuals from the Christchurch Health and Development Study (CHDS) which evaluated differentialDNA methylation in response to regular cannabis use [12]. The CHDS is a longitudinal study of a birth cohort of 1265 children born in 1977 in Christchurch, New Zealand, who have been studied on 24 occasions from birth to the age of 40 (n = 904 at age 40). Of this, a total 96 individuals were selected, and arrays were performed in two separate batches in consecutive years (n = 48 per year).

For validation analysis we selected individuals with EPIC array data, as well as new individuals (n=82), to serve as a validation and expansion cohort for the differential DNA methylation identified via EPIC array [12]. Specifically, we asked whether BSAS, after determination of the most appropriate normalisation method, produced the same average methylation values as EPIC arrays, when comparing case data to control data.

Interestingly, while both EPIC array and BSAS are readily used as standalone experiments, direct comparison between the two technologies is not widely reported. Given the rising popularity of studies investigating DNA methylation, establishing the reliability of a method that would allow for accurate expansion of existing studies in larger cohorts would be valuable to the scientific community.

## Methods and materials

### Cohort selection and DNA extraction - EPIC arrays

EPIC array data used in this study has previously been published[12]. Briefly, in this study we use DNA from human participants who are partitioned into three groups: i) regular cannabis users, who had never used tobacco (“cannabis-only”); those who consumed both cannabis and tobacco (“cannabis plus tobacco”), and; iii) controls, who consumed neither cannabis nor tobacco. Controls were matched as closely as possible for sex, ethnicity and parental socioeconomic status (data described in [12]). DNA was extracted from whole blood using the KingFisher Flex System (Thermo Scientific, Waltham, MA USA), as per the published protocols. DNA was quantified via NanoDrop™ (Thermo Scientific, Waltham, MA USA) and standardised to 100ng/μl. Equimolar amounts were shipped to the Australian Genomics Research Facility (AGRF, Melbourne, VIC, Australia) for processing via the Infinium® Methylation EPIC BeadChip (Illumina, San Diego, CA USA).

### Bioinformatic analysis – processing and normalisation of raw EPIC array data

For this study, analysis was carried out using R statistical software (Version 3.5.2)[33]. Quality control was first performed on the raw data; sex chromosomes and 150 failed probes (detection P value greater than 0.01 in at least 50% of samples) were excluded from analysis. Furthermore, potentially problematic CpGs with adjacent single nucleotide polymorphisms (SNPs), or that did not map to a unique location in the genome [34] were also excluded. The raw data were then normalised using four different pipelines, Illumina, SWAN [35], Funnorm [36] and Noob [37] in the minfi package [38]. Normalisation was then checked by observing density plots as well as multidimensional scaling plots of the 5000 most variable CpG sites.

### Cohort selection and DNA extraction – BSAS experiments

BSAS analysis was carried out on two groups: cannabis plus tobacco users (n=44) and controls (n=38), who had never used cannabis – in contrast to the EPIC array analysis, no cannabis-only participants were used in BSAS. This is a consequence of the small number of individuals who use cannabis but who do not also use tobacco. Cannabis users were all selected on the basis that they either met DSM-IV diagnostic criteria [39] for cannabis dependence or had reported using cannabis consumption on a daily basis for a minimum of three years prior to age 28. Participants were matched as closely as possible for sex, ethnicity, and parental socioeconomic status (Supplementary Table 1). Because this was a birth cohort collected across a four month period, they are all of a similar age. Collection and analysis of DNA in the Christchurch Health and Development Study was approved by Southern Health and Disability Ethics Committee (CTB/04/11/234/AM10). DNA was extracted from whole blood samples using a Kingfisher Flex System (Thermo Scientific, Waltham, MA USA). DNA was quantified via nanodrop (Thermo Scientific, Waltham, MA USA) and standardised to 100 ng/μl. Bisulfite treatment was carried out using the EZ DNA Methylation-Gold kit (Zymo Research, USA) as per the manufacturer’s instructions. DNA samples were then diluted to a final concentration of 100 ng/μl.

**Table 1:** CpG site differences from EPIC array and the BSAS methods at the 15 loci of differing levels of significance (not significant, nominally significant, significant after P-value adjustment)

|  |  |  | Illumina EPIC array |  |  | BSAS |  |  | Difference between methods |
| --- | --- | --- | --- | --- | --- | --- | --- | --- | --- |
| Cg/Gene | Position in genome | Illumina ID | $\beta$ difference | P value | FDR Adjusted P value | $\beta$ difference | P value | FDR Adjusted P value | $\beta$ difference |
| 1 *AHRR2* | Chr5, GB | cg05575921 | -0.233 | 5.33E-12 | 3.7E-06 | -0.041 | 0.006* | 0.245 | -0.192 |
| 2 cg11977356* | Chr19 | cg11977356 | -0.040 | 0.474 | 0.999 | -0.004 | 0.406 | 0.959 | -0.036 |
| 3 *ITPR1* | Chr3, GB | cg08987995 | -0.001 | 0.572 | 0.999 | 0.005 | 0.820 | 0.822 | -0.006 |
| 4 *MAGI* | Chr7, GB | cg21121803 | -0.008 | 0.572 | 0.999 | -0.007 | 0.809 | 0.959 | -0.0004 |
| 5 *EHMT2* | Chr6, GB | cg07829740 | 0.005 | 0.037 | 0.999 | -0.015 | 0.071 | 0.579 | 0.020 |
| 6 *PPM1L* | Chr3, GB | cg26406186 | -0.006 | 0.818 | 0.999 | 0.011 | 0.904 | 0.963 | -0.017 |
| 7 cg00571101* | Chr12 | cg00571101 | 0.004 | 0.368 | 0.999 | -0.004 | 0.813 | 0.952 | 0.008 |
| 8 cg09078959* | Chr5 | cg09078959 | -0.001 | 0.893 | 0.999 | -0.005 | 0.001* | 0.245 | 0.004 |
| 9 cg01614625* | Chr7 | cg01614625 | -0.009 | 0.370 | 0.999 | -0.006 | 0.569 | 0.952 | -0.004 |
| 10 *DP10* | Chr2, GB | cg05868547 | 0.006 | 0.077 | 0.999 | -0.003 | 0.713 | 0.952 | 0.009 |
| 11 cg11293828* | Chr12 | cg11293828 | -0.014 | 0.665 | 0.999 | 0.032 | 0.735 | 0.952 | -0.045 |
| 12 *CHD7* | Chr5, 5'UTR | cg19926587 | -0.007 | 0.960 | 0.999 | -0.006 | 0.429 | 0.959 | -0.001 |
| 13 *NIPAL4* | Chr5, TSS1500 | cg17695979 | -0.007 | 0.714 | 0.999 | -0.003 | 0.106 | 0.713 | -0.004 |
| 14 *PRDM5* | Chr4, GB | cg01118724 | -0.004 | 0.734 | 0.999 | 0.005 | 0.116 | 0.713 | -0.009 |
| 15 *SLC17A7* | Chr19, GB | cg02624701 | -0.043 | 0.312 | 0.999 | 0.018 | 0.646 | 0.952 | -0.061 |

### CpG site selection, primer design and amplification - BSAS

A total of 15 CpG sites, representing 15 individual probes from the Illumina EPIC array were chosen based on their differential methylation status in cannabis plus tobacco users compared to controls (Table 1). A range of probes at differing levels of significance (not significant, nominally significant, significant after P-value adjustment) were chosen to reflect the range of data provided by the EPIC arrays. Primers to amplify bisulfite-treated DNA were designed using the online tool BiSearch [40] to amplify a ~250 base pair region which spanned the CpG site (Supplementary Table 2). At the 5’ end of each primer sequence, an Illumina overhang (33 base pair sequence) was included to ensure the ability to pool the amplicons and barcode them for high-throughput sequencing.

Bisulfite-converted DNA was amplified via PCR, using KAPA Taq HotStart DNA Polymerase (Sigma, Aldrich) under the following conditions: 95 °C for 10 min, followed by 40 cycles of 95 °C for 30 sec, 59 °C for 20 sec, 72 °C for 7 min, and finally held at 4 C° using the Mastercycler Nexus (Eppendorf, Australia). PCR products were then purified with the Zymo DNA Clean & Concentrator Kit™ (Zymo Research, USA).

Following the PCR, DNA was cleaned up with Agencourt® AMPure® XP beads (Beckman Coulter) and washed with 80% ethanol and allowed to air-dry. DNA was then eluted with 52.5 μl of 10 mM Tris pH 8.5 before being placed back into the magnetic stand. Once the supernatant had cleared, 50 μl of supernatant was taken up and aliquotted into a fresh 96-well plate. DNA samples were quantified using the Quant-iT™ PicoGreen™ dsDNA Assay kit (Thermo Fisher) using the FLUROstar® Omega (BMG Labtech). Sequence libraries were prepared using the Illumina MiSeq™ 500 cycle Kit V2, and sequenced on an Illumina MiSeq™ system at Massey Genome Services (Palmerston North, New Zealand).

### Bioinformatic and statistical analysis – BSAS data

Illumina MiSeq™ sequences were trimmed using SolexQA++ software and aligned to FASTA bisulfite converted reference sequences using the package Bowtie2 (version 2.3.4.3). Each individual read was then aligned to all reference sequences using the methylation-specific package Bismark [41]. Bismark produced aligned mapped reads with counts for methylated and unmethylated cytosines at each CpG site, thus BSAS returns additional CpG sites to the intended validation target, as each sequencing read contains multiple CpG sites. Cytosine proportion is calculated based upon the number of cytosines divided by the number of cytosines with the additions of the number of thymines present(C/(C_l_)+ T). This gave the average methylation β values for each individual at each given CpG site. These β values could be anywhere between 0 −1, with a β equal to 1 indicating 100% methylation at that CpG site across all sequencing reads. These data were imported into R Studio (RStudio version 3.3.0) and the edgeR package [42] was used to assess differential DNA methylation between cannabis users and controls; coverage level was set to greater or equal to “8” across unmethylated and methylated counts. This was also set at 50 and 100 reads and no differences were seen between the results at any of these thresholds, so “8” was used for the continuation of BSAS calling under the recommendations of [42]. A negative binomial generalised model is used to fit the counts (methylated and unmethylated reads) in regards to the two variable groups.

A scatter plot including a linear regression line with adjusted R^2^ values was generated in R Studio to quantify the correlation between β values produced with EPIC array and BSAS. Adjusted R^2^ values were calculated for: i) BSAS cases versus EPIC cases, and; ii) BSAS controls versus EPIC controls. A Bland Altman analysis [43] was used to compare the agreement of the two techniques. Means were log transformed and lower and upper levels of agreement with 95% confidence intervals were calculated. Summary tables compiled of the CpG sites of interest with nominal P value significance and post multiple testing using false discovery rate (FDR) of less than 0.05 were considered to be statistically significant. All graphs were constructed using the R package ggplot2 [44].

## Results

### Validation and replication of EPIC array data using BSAS

The differences between β values of cannabis plus tobacco users (cases) and controls were calculated for each method (EPIC array and BSAS, Table 1). All sites except for cg05575921 in *AHRR2* demonstrated DNA methylation level differences between analysis platforms of less than 6%.

Correlations were plotted individually for cases and controls for the two detection methods. BSAS versus EPIC cases resulted in an adjusted R^2^ of 0.8878 and BSAS versus EPIC controls gave an adjusted R^2^ of 0.8683 (Fig 1).

**Figure 1.**
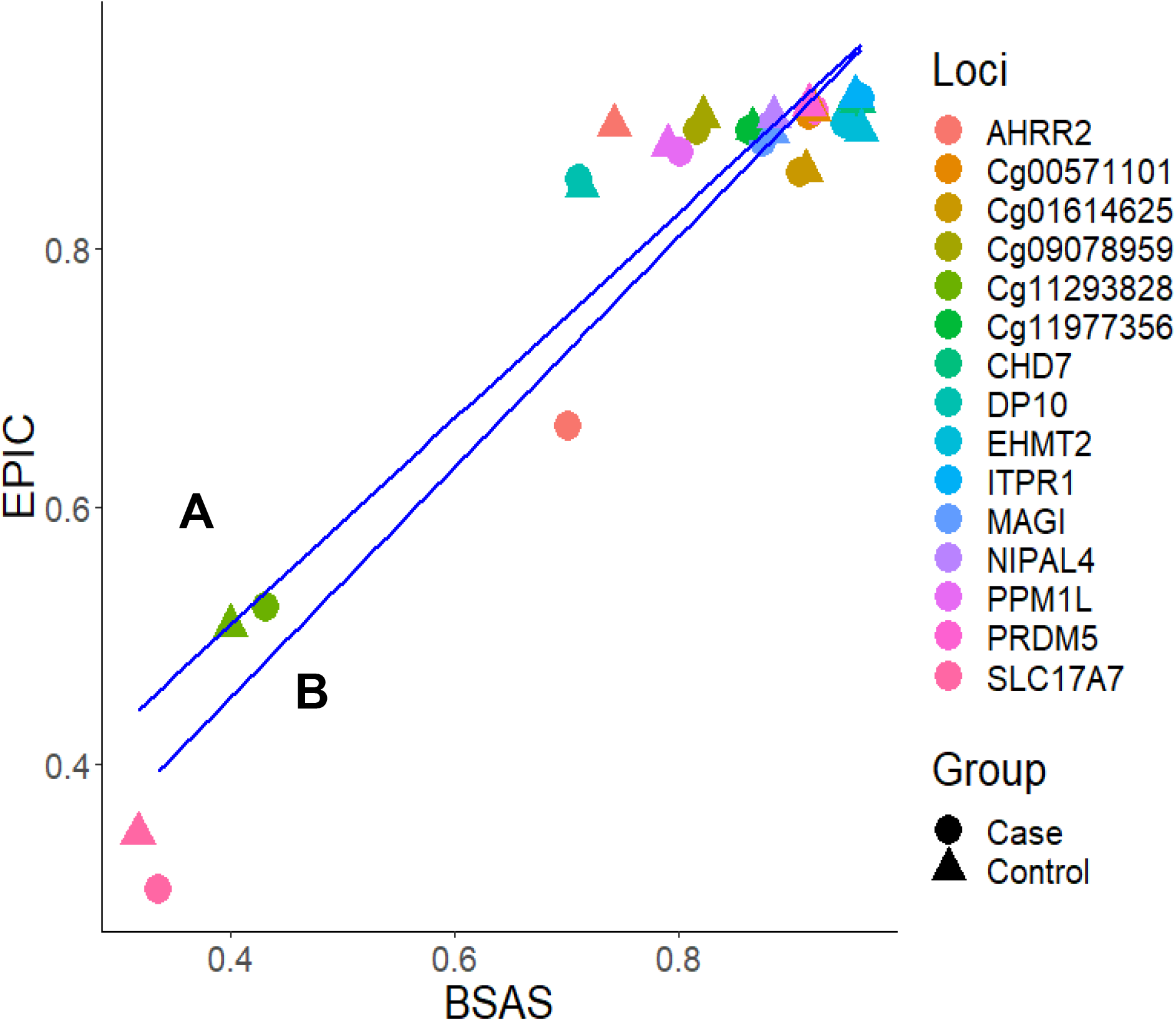
Scatter plot with a linear regression of the β values at each locus for BSAS and EPIC array plotted against each other. Colours represent the loci of interest, with the shapes representing the case and controls. There are two regression lines: A represents the correlation between cases with an adjusted R^2^ = 0.8878 and B represents controls with R^2^ = 0.8683.

### Do these two methods of DNA methylation detection correlate?

A Bland Altman analysis was carried out on the loci investigated by BSAS and compared to data for the same loci produced using the Illumina EPIC array. Fig 2A shows cannabis users (cases) measured using BSAS and the EPIC array on the X axis, while the Y axis represents the differences between the measurements. The observed differences between loci in cannabis cases (EPIC and BSAS) fall within the lines of agreement. Fig 2B shows the control group differences plotted for the same loci for BSAS and the EPIC array methods. Similar to above, all data points fall within the lower and upper lines of agreement.

**Figure 2.**
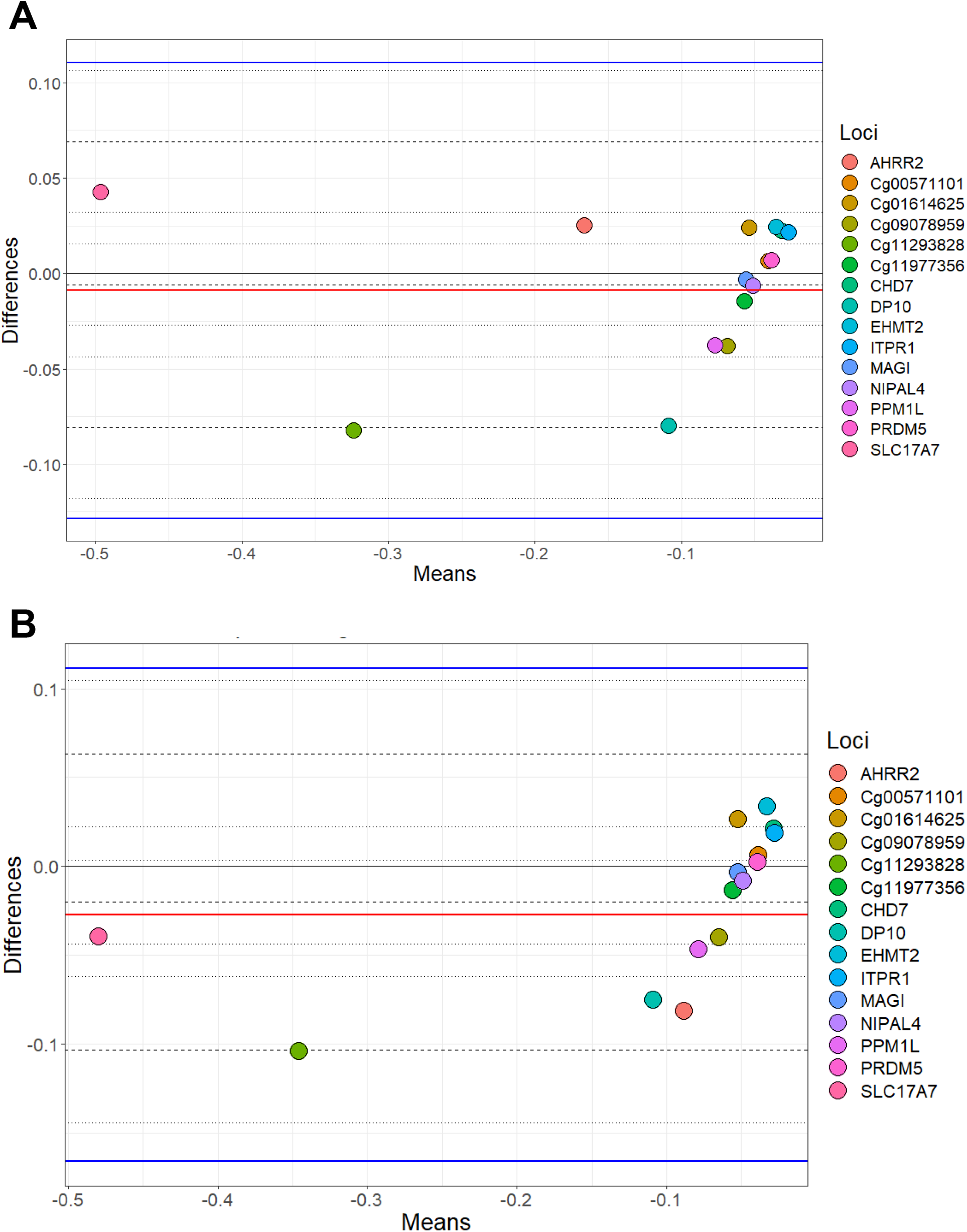
Bland Altman plots showing the mean differences between DNA methylation as measured by EPIC array vs. the same CpG sites measured using BSAS. A) Data from cannabis users, gathered using BSAS and the EPIC array (Cases) B) the control subjects used in BSAS and EPIC array. Each of the 15 points represent the CpG sites investigated.

## Discussion

High throughput array technologies have facilitated the next step in assessing associations between DNA methylation changes in response to a known environmental exposure at a genome-wide level. The EPIC array (as well as the predecessor 27k and 450k arrays) is one such platform that allows for the characterisation of these DNA methylation changes. Through these approaches, various studies have furthered our understanding of how DNA methylation can play a role in response to different environmental exposures.

We selected the orthogonal method BSAS to determine its applicability as a validation, replication and/or expansion tool for EPIC array. BSAS is often used as a standalone method for assessing differential DNA methylation at specific CpG sites, usually because it is more cost-effective than EPIC arrays, and allows analysis of many samples at once, in multiplex. It returns data for all CpGs within a targeted region of interest (~250 base pairs) with results providing base pair-level specificity [31]. In this study, BSAS estimation of differential DNA methylation correlated with differential methylation determined via EPIC array. However, although the data correlates between the methods (adjusted R^2^ cases, 0.8878 and adjusted R^2^ controls, 0.8683), we urge caution when interpreting this correlation as proof that BSAS will be a suitable independent validation of EPIC array data in every experiment. This is because while the data presented here correlated between BSAS and EPIC array as a whole dataset, some sites showed larger differences between average methylation estimated using BSAS vs. EPIC array. In particular, where the differential methylation on EPIC array was greater than 5% between cases and controls, BSAS was unable to detect this differential DNA methylation to the same magnitude as EPIC array. For instance, *AHRR2* exhibited a 4% difference in methylation between cases and controls when assessed using BSAS (the highest value detected in using BSAS in this study), compared to 23% using EPIC array. Thus, while a strong correlation between EPIC array data and BSAS data was found across the 15 CpG sites investigated, which itself implies an association between the average methylation at each CpG for the two techniques, further work on CpG sites with higher magnitude changes is needed to determine whether BSAS is limited by the magnitude of differential methylation it is able to detect. However, it is worth noting that most studies of differential methylation report modest (<5%) significant differential methylation observations, suggesting that BSAS may prove useful, given inclusion of rigorous controls of known differential methylation to ensure accuracy of results.

Due to the sequence-based nature of BSAS data (compared to the probe-based nature of EPIC arrays) BSAS, as a standalone method, offers some advantages that are not applicable to EPIC arrays. For instance, BSAS has the potential to determine novel differentially methylated CpGs which may be near (in the same targeted region) but not the initial pre-determined CpG site of interest. This is possible because all CpGs within an e.g. 500 base pair region are returned using BSAS data, only one of which may be on an EPIC array. Further, via this targeted sequencing process, BSAS may reveal novel differentially methylated regions (DMRs). DMRs are described as areas which exhibit multiple successive methylated CpG sites which may have biological impact within the genome. For example, they have been implicated in the development and progression of disease [45]. Therefore targeting more than a single CpG site may provide further insight into genes and regions of interest. Consequently, while here we have used BSAS technology to replicate/validate differential methylation identified via EPIC array, given that BSAS outputs largely correlate with EPIC data, equally, BSAS could be as a “discovery-based tool”; if significantly differentially methylated CpGs are identified via BSAS, this would serve to justify further investigation using a robust and more expensive high throughout method.

The EPIC array still remains the most reproducible way to measure DNA methylation [46]. This is because the probe-based nature of the method frequently produces comparable results across research groups and arrays. For example, detection of differential methylation using the EPIC array found a difference of 23% in cannabis plus tobacco users, compared to controls, at *AHRR2* (cg05575921, Table 1), a result that is supported by other studies in tobacco smokers using EPIC array [9, 47–50]. *AHRR2* has an important role in controlling a range of different physiological functions; it contributes to regulation of cell growth, regulation of apoptosis and contributes to vascular and immune responses [51–54].

BSAS and EPIC array rely upon different chemistries and methods to detect DNA methylation. This may account for the majority of the variation found between the two methods. BSAS relies upon PCR amplification of DNA that is treated with sodium bisulfite. When DNA is treated, unmethylated cytosine residues are converted into uracils via hydrolytic deamination. Amplification of these uracil nucleotides during this process are replaced by thymines during replication and the 5-methylcytosines are left unreactive throughout the deamination process and then are amplified as cytosines. It then becomes possible to ‘read’ values of methylation for each cytosine in an amplicon via DNA sequencing [55]. The ability to treat DNA with sodium bisulfite has led to the expansion of research undertaken within this field [56]. However, it is important that we ensure the validity of the results are not limited by the manner in which the data was produced. Ensuring that we limit these discrepancies between technologies will allow for better validation of data. There is potential for errors to occur at this step, because incomplete bisulfite conversion cannot be distinguished from 5-methylcytosine, this can possibly introduce false positive methylation calls at this point [57] [58]. Although both techniques rely upon bisulfite treatment, it is this source of error followed by the PCR amplification that might explain the differences in results we have observed. Refining these sources of error may provide much more comparable results between the two methods.

In conclusion, we chose to validate EPIC array data by using the alternative method, BSAS, to detect differential methylation at CpG sites. However, it is possible that BSAS may be unable to reproduce the magnitude of changes that are shown in the EPIC array system, which may be a consequence of lack of specificity and addition error rate through PCR amplification. It does however, have the ability to assess differentially methylated regions, rather than individual CpG sites. As some regions of the genome are more susceptible to methylation change than others, BSAS could detect swathes of correlated differential methylation at neighbouring CpG sites in certain areas of the genome. From the results shown here, BSAS has the potential to be able to detect methylation marks such as metastable epialleles, which maybe hallmarks for disease later on in life. Finally, although BSAS does not generate the same significance level as the EPIC array in some instances, we demonstrate that BSAS can be used as an investigative tool for specific regions of the genome.

## Acknowledgments

University of Otago Research Grant to M.A.K., The Carney Centre for Pharmacogenomics. CHDS was funded by the Health Research Council of New Zealand (Programme Grant 16/600) and the Canterbury Medical Research Foundation.

